# Bidirectional pharmacological perturbations of the noradrenergic system differentially affect tactile detection

**DOI:** 10.1101/2020.04.22.052043

**Authors:** Jim McBurney-Lin, Yina Sun, Lucas S. Tortorelli, Quynh Anh Nguyen, Sachiko Haga-Yamanaka, Hongdian Yang

## Abstract

The brain neuromodulatory systems heavily influence behavioral and cognitive processes. Previous work has shown that norepinephrine (NE), a classic neuromodulator mainly derived from the locus coeruleus (LC), enhances neuronal responses to sensory stimuli. However, the role of the LC-NE system in modulating perceptual task performance is not well understood. In addition, systemic perturbation of NE signaling has often been proposed to specifically target the LC in functional studies, yet the assumption that localized (specific) and systemic (nonspecific) perturbations of LC-NE have the same behavioral impact remains largely untested. In this study, we trained mice to perform a head-fixed, quantitative tactile detection task, and administered an α2 adrenergic receptor agonist or antagonist to pharmacologically down- or up-regulate LC-NE activity, respectively. We addressed the outstanding question of how bidirectional perturbations of LC-NE activity affect tactile detection, and tested whether localized and systemic drug treatments exert the same behavioral effects. We found that both localized and systemic suppression of LC-NE impaired tactile detection by reducing motivation. Surprisingly, while locally activating LC-NE enabled mice to perform in a near-optimal regime, systemic activation impaired behavior by promoting impulsivity. Our results demonstrate that localized silencing and activation of LC-NE differentially affect tactile detection, and that localized and systemic NE activation induce distinct behavioral changes.

## 1. Introduction

The locus coeruleus (LC) is a major source of the neuromodulator norepinephrine (NE) in mammalian brains. With profuse projections across the central nervous system, this modulatory circuit has been hypothesized to be critical in mediating a variety of brain functions and behavior, including sleep-wake transition, perception, attention and learning. The dysfunction of the LC-NE circuit has also been thought to be involved in several neurological disorders (Arnsten, 2000; Berridge and Waterhouse, 2003; Aston-Jones and Cohen, 2005; Sara, 2009; Sara and Bouret, 2012; Waterhouse and Navarra, 2019).

We recently proposed that understanding how LC-NE modulates sensory perception offers a stepping stone toward unraveling its roles in higher cognitive functions (McBurney-Lin et al., 2019). LC neurons extensively innervate sensory cortical and subcortical regions, and LC-NE signaling modulates sensory neuron responses to external stimuli (e.g., (Foote et al., 1975; Waterhouse et al., 1980; Kasamatsu and Heggelund, 1982; Morrison and Foote, 1986; Simpson et al., 1997; Devilbiss and Waterhouse, 2004; Manella et al., 2017; Navarra et al., 2017; Rho et al., 2018)). LC-NE may also affect sensory perception through modulating motivation or attention (Berridge and Waterhouse, 2003; Lee and Dan, 2012; Sara and Bouret, 2012; Thiele and Bellgrove, 2018). To our knowledge, only a limited number of studies have examined LC-NE influence on perception-related behavior (Doucette et al., 2007; Escanilla et al., 2010; Martins and Froemke, 2015; Navarra et al., 2017; Rodenkirch et al., 2019). It remains poorly understood how bidirectional perturbations of LC-NE activity affect perceptual task performance.

To examine the causal role of a neural circuit, such as the LC-NE, in regulating behavior, one would perturb this system and assess the subsequent behavioral changes. Traditional lesion approaches may induce compensatory plasticity changes (Acheson et al., 1980; Harik et al., 1981; Valentini et al., 2004) and mask the effects specific to LC-NE. More recent studies employed acute, reversible perturbations including pharmacological, electrical, chemogenetic, and optogenetic stimulations. Among these approaches, pharmacology facilitates translational comparison between animal and human studies. The inhibitory α2 adrenergic receptors (ARs) are highly expressed in the LC, but only sparsely expressed, if at all, in neighboring brainstem regions (McCune et al., 1993; Nicholas et al., 1993). Targeting α2 ARs is considered a specific manner to perturb LC-NE activity (e.g., (Neves et al., 2018)). Agonizing α2 ARs suppresses LC-NE signaling by hyperpolarizing LC neurons and reducing NE release in downstream areas (Cedarbaum and Aghajanian, 1977; Aghajanian and VanderMaelen, 1982; Abercrombie and Jacobs, 1987; Aghajanian and Wang, 1987; Adams and Foote, 1988; Berridge et al., 1993; Kalwani et al., 2014). Conversely, antagonists acting on α2 ARs increase LC neuron excitability and spiking response to stimuli as well as NE release (Cedarbaum and Aghajanian, 1976; Aghajanian and VanderMaelen, 1982; Raiteri et al., 1983; Rasmussen and Jacobs, 1986; Simson and Weiss, 1987; Adams and Foote, 1988; Herr et al., 2012).

Human studies have reported that systemically up- or down-regulating NE signaling (mainly through targeting α2 ARs) affected subjects performing perception-related tasks (Halliday et al., 1989; Turetsky and Fein, 2002; Gelbard-Sagiv et al., 2018). Targeting α2 ARs nonspecifically (e.g., intraperitoneal - i.p. or intracerebroventricular - i.c.v., hereafter referred to as “systemic”) or specifically (e.g., intra- or peri-LC, hereafter referred to as “localized”) exerts similar changes on LC activity (Aghajanian and VanderMaelen, 1982; Adams and Foote, 1988; Berridge et al., 1993). However, systemic perturbations of α2-ARs could induce physiological and behavioral effects that are different from localized perturbation. Systemic α2 perturbation would likely affect noradrenergic neurons in the nucleus of the solitary tract (Van Bockstaele et al., 1999; Kirouac, 2015), as well as many α2-expressing regions in the nervous system (McCune et al., 1993; Nicholas et al., 1993; Robertson et al., 2013). It should also be noted that α2-ARs are expressed both presynaptically (auto-receptors) and postsynaptically in terminal fields. Agonizing or antagonizing presynaptic α2-ARs suppresses or enhances NE release, respectively, and the postsynaptic effects would depend on the specific types of postsynaptic adrenergic receptors that are activated in terminal fields. In contrast, agonizing or antagonizing postsynaptic α2-ARs exerts direct inhibitory or excitatory postsynaptic effects, respectively.

Head-fixed behavior facilitates stimulus control and movement measurement, and allows reliable quantification of different components of perceptual behavior, including detection, discrimination, impulsivity and motivation (Schwarz et al., 2010; Guo et al., 2014). To our knowledge, using well-controlled, quantitative perceptual behavior to examine the effects of localized (specific) and systemic (nonspecific) perturbations of LC-NE is lacking.

In the current study, we trained mice to perform a head-fixed, quantitative tactile detection task. We administered an α2 agonist or antagonist to pharmacologically down- or up-regulate LC activity, respectively. We addressed the outstanding question of how bidirectional perturbations of LC activity affect tactile detection, and tested whether localized and systemic drug treatments exert the same behavioral effects.

## 2. Methods

### 2.1 Mice

Both male and female mice were used in this study. All mice were C57BL/6J except 2 (out of 6) included in the localized clonidine treatment were of mixed B6J/129 background. Mice were housed with reversed light/dark cycle (9A - 9P dark, 9P - 9A light). Mice of 6-12 weeks were implanted with head posts and/or cannulae. Clonidine (an α2 agonist, Sigma-Aldrich) was administered locally in 6 mice and systemically in 3 mice. Yohimbine (an α2 antagonist, Sigma-Aldrich) was administered locally in 7 mice and systemically in 5 mice. Every mouse received corresponding localized or systemic saline injections as controls. Quantification of localized pharmacological effects on LC activity was performed by immunostaining for the immediate early gene c-fos in a separate group of 11 mice. All procedures were approved by the UC Riverside Animal Care and Use Committee.

### 2.2 Surgery

Head post surgeries were similar to previously published work (Yang et al., 2016). In brief, mice were anesthetized (1-2% isoflurane) and affixed to a stereotaxic instrument (Kopf, RWD). Body temperature was maintained with a heating blanket (Harvard Apparatus, RWD) throughout the surgical procedures. The scalp over the dorsal surface of the skull was cleaned with betadine and 70% ethanol, and removed. The periosteum was removed and the skull scored with a dental drill. Cyanoacrylate was applied to the border of the skull and scalp. The head post was placed and secured with dental acrylic. A craniotomy of ~1 mm × 1 mm was made over the left hemisphere, centered at 5.2 - 5.3 mm posterior to bregma and 0.9 - 1.0 mm lateral to midline. A guide cannula (27G, 3.5 mm long, RWD) with dummy insert was advanced vertically into the brain until a depth of 1.8 mm. Dental acrylic was used to secure the guide cannula and filled in the remaining exposed skull surface. After surgery, mice were single housed and allowed to recover for at least 48 hours.

### 2.3 Behavioral Task

Following recovery from the surgery, mice were restricted to 1 mL/day water consumption for 7-10 days before behavioral training. The behavior task was adapted from published work (Yang et al., 2016). Briefly, mice were trained to perform a head-fixed, Go/NoGo single-whisker detection task, in which mice reported whether they perceived a brief deflection (200 ms, 25 Hz, ~600 deg/s) to the right C2 whisker by licking or withholding licking. Ambient white noise (1 - 40 kHz) was played throughout the session. An auditory cue (8 kHz) was presented at the beginning of each trial, 1.5 s prior to the time of possible stimulus onset. Trial outcomes comprised a mixture of successful and failed stimulus detection (Hit and Miss), as well as successful and failed responses to stimulus absence (Correct Rejection and False Alarm). Trials were aborted if mice licked prematurely during the waiting period between auditory cue and the time of possible stimulus onset (Impulsive). Trials were also considered impulsive when mice licked within the first 100 ms window from stimulus onset (Mayrhofer et al., 2013; Yang et al., 2016). Mice performed one behavior session (300-500 trials) per day. Mice never achieved saturating performance in this task (Yang et al., 2016), indicating that detecting weak single-whisker deflection is perceptually demanding. All aspects of behavioral control were managed by custom Arduino-based hardware and software. Behavioral data were acquired with WaveSurfer (https://www.janelia.org/open-science/wavesurfer).

### 2.4 Pharmacology

All drugs were dissolved in physiological saline. Localized pharmacology was administered during behavior sessions. Drug or saline was loaded into a 1 μL Hamilton syringe, controlled by a syringe pump (Harvard Apparatus). Mice were placed in the behavior chamber, and injection cannula (33G, 5 mm long) inserted into the guide cannula. The infusion depth was 3.3 mm. Infusion was initiated within the first 20 behavior trials. 300 nL of drug or saline was infused at a rate of 60 nL/min. At the conclusion of a behavior session, injection cannula was removed and dummy insert replaced.

Systemic pharmacology was administered just prior to behavior sessions. Mice were briefly anesthetized (< 1 minute) with 2-3% isoflurane, during which 50 μL of drug or saline was injected via i.p.. Mice were allowed to recover for 5 minutes before starting the behavior session. During baseline behavioral sessions (one day before i.p. treatment), mice were also briefly anesthetized to account for any potential effects from anesthesia.

### 2.5 Histology

At the conclusion of behavioral experiments, mice with cannula implants received localized Fluoro-Gold infusion (0.1-1%, 300 nL) at a rate of 60 nL/min. 40-60 minutes later, mice were anesthetized with ketamine/xylazine and perfused intracardially with 4% paraformaldehyde, and the brains harvested and post fixed. 100 μM thick coronal sections were cut (Leica, VT1200s). Sections containing LC were incubated with rabbit anti-Tyrosine Hydroxylase (TH) antibody (Thermofisher OPA 1-04050, 1:1000), followed by goat anti-rabbit IgG Alexa Fluor 488 or 594 secondary antibody (Thermofisher A32731 or A32740, 1:1000), and mounted with DAPI mounting media (Vector labs). Co-localization of Fluoro-Gold and TH immunoactivity, as well as the cannula tract, were used to verify cannula placement.

The expression of an immediate early gene, c-fos, was examined to assess the impact of localized drug treatment on LC activity. Infusions were performed in the left LC, with the contralateral (right) LC serving as a control. Clonidine was infused in 4 awake mice. Yohimbine was infused in 5 mice, 2 of which received infusion under anesthesia, with the purpose to reduce basal LC activation and enhance the contrast between the injected side and the control side. The remaining 3 mice received infusion during wakefulness. Saline was infused in 2 awake mice. All mice were perfused 40-60 minutes post infusion. Coronal sections containing LC were first incubated with rabbit anti-c-fos antibody (Cell Signaling 2250S, 1:400), followed by secondary antibody (Thermofisher A32740, 1:400). Sections were then incubated with rabbit IgG isotype control (Thermofisher 31235, 1:17000) to quench nonspecific signals, and subsequently stained for TH. Z-stack images were acquired using a confocal microscope (Leica SPE II) and flattened using Fiji (Schindelin et al., 2012).

### 2.6 Data Analysis

Behavior data were analyzed off-line with MATLAB. To account for the fact that some mice did not immediately engage in the task, the initial 20-40 trials were removed from behavior analysis. In some sessions, trials toward session end were also removed from analysis when mice appeared to be disengaged from the task (Hit rate dropped below 50%, typically after 300-400 trials). For sessions shown in Fig. 1, we included an additional 20-50 trials toward session end to demonstrate a near-complete cessation of task performance. Decision bias/criterion (c) and detection sensitivity (d’) were calculated based on Hit rate (HR) and False Alarm rate (FAR): *c = z(HR) - z(FAR), d’ = −(z(HR) + z(FAR))/2*, where z is the normal inverse cumulative distribution function.

**Figure 1.**
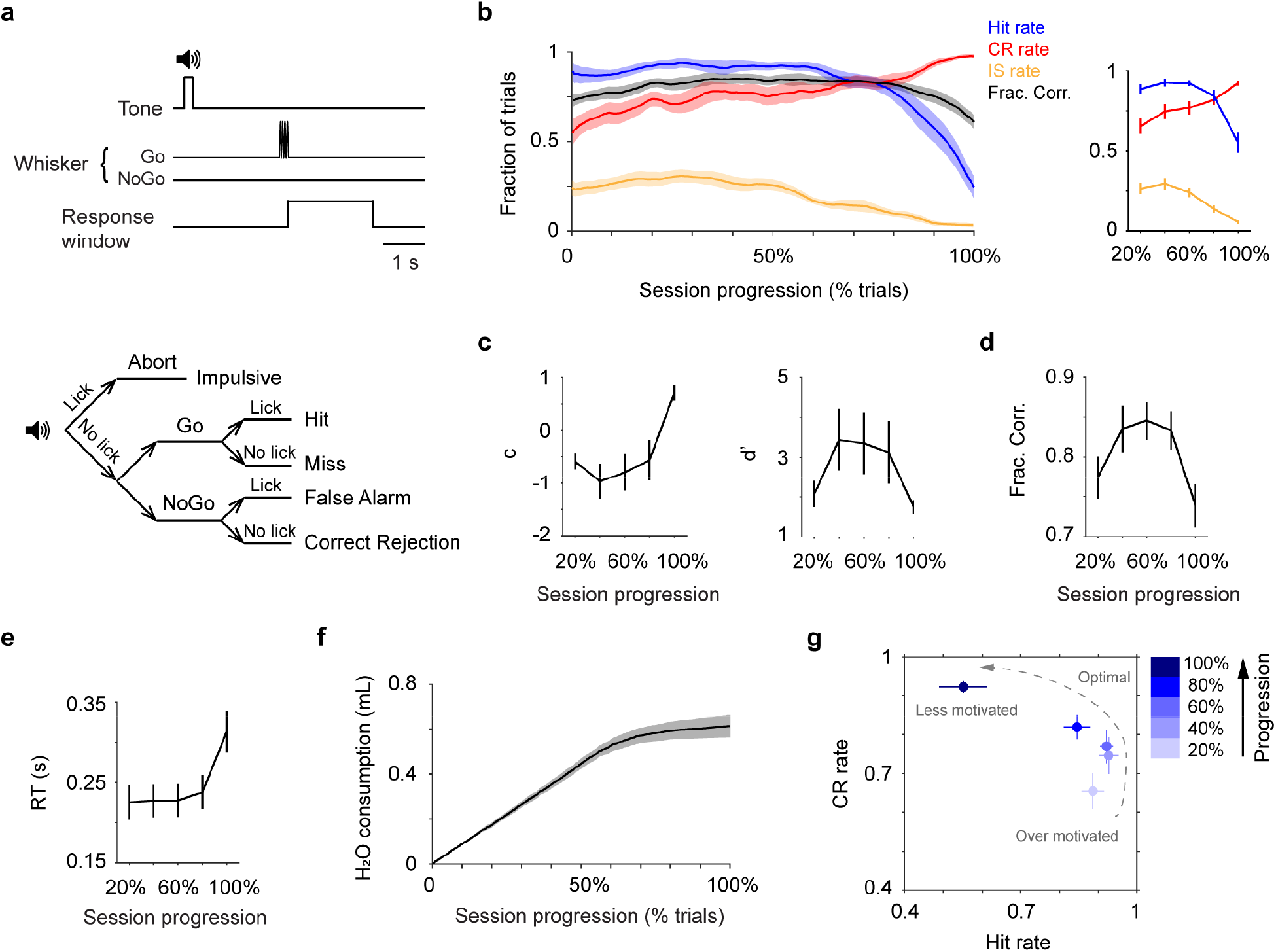
Mouse behavior fluctuates within single sessions. **a.** Trial structure (top) and the five possible trial types (bottom). **b.** Left: Mean single-session trajectories of Hit rate, CR rate, IS rate and overall performance (± s.e.m.). Behavioral sessions of different lengths (348 ± 20 trials, mean ± s.e.m., n = 13) are normalized using % total number of trials (session progression). Trajectories are smoothed using a moving window of 30 trials. Right: Trajectories of Hit, CR and IS rates averaged every 20% progression. **c.** Trajectories of decision bias (c) and detection sensitivity (d’), based on Hit and CR rates in **b**. **d.** Trajectory of overall performance (Fraction Correct shown in **b**, averaged every 20% progression) illustrates an inverted-U shape. **e**. Mean single-session trajectory of RT (± s.e.m.), averaged every 20% progression. **f.** Mean single-session trajectory of cumulated water consumption (± s.e.m.), based on an estimate of 5 μL dispense per Hit trial. **g.** CR rate vs. Hit rate trajectory, based on values in **b**. CR, Correct Rejection; IS, Impulsive; RT, Reaction Time.

c-fos expression was analyzed using QuPath (Bankhead et al., 2017). Borders around the LC were manually drawn to identify regions of interest. For each mouse, 2-3 images with the greatest TH and c-fos expressions were used to determine the minimum and maximum cell sizes, as well as the fluorescent intensity threshold. Individual cells expressing supra-threshold TH or c-fos were detected. Results were manually verified for each image.

Data were reported as mean ± s.e.m. unless otherwise noted. Statistical tests were by two-tailed Wilcoxon signed rank unless otherwise noted.

## 3. Results

### 3.1 Mouse behavior fluctuates within single sessions

Mice were trained to perform a head-fixed, Go/NoGo single-whisker detection task, in which mice reported whether they perceived a brief deflection to the right C2 whisker by licking or withholding licking (Fig. 1a). The performance of well-trained mice fluctuated during single behavior sessions, as reported by others recently (Berditchevskaia et al., 2016). A typical behavior session started with mice licking indiscriminately, resulting in high Hit rate (fraction of Hit trials among Go trials), high Impulsive rate (IS rate, fraction of IS trials among all trials), and low Correct Rejection rate (CR rate, fraction of CR trials among NoGo trials). As the session proceeded, Hit rate remained high while mice better withheld licking in NoGo trials, increasing Correct Rejection rate. Towards session end, mice licked less in all trials, and Hit and Impulsive rates reached a minimum and Correct Rejection rate reached a maximum (Fig. 1b). Within sessions, the fluctuations of Impulsive rate were positively correlated with Hit rate, and highly anti-correlated with Correct Rejection rate (Fig. S1). Using signal detection theory (Green and Swets, 1966), we found that decision bias/criterion (c) increased over time, while detection sensitivity/discriminability (d’) exhibited an inverted-U profile (Fig. 1c). Toward session end, reaction time (RT, latency from stimulus onset to the time of first licking response) increased and lick frequency declined (Fig. 1e, Fig. S2). As demonstrated in previous work (e.g., (Dickinson and Balleine, 1994; Mayrhofer et al., 2013; Berditchevskaia et al., 2016)), these behavioral changes reflect a systematic shift of the motivational states of the mice. To illustrate this shift, we constructed a trajectory of motivational states based on Hit rate and Correct Rejection rate (Fig. 1g): mice started with an over-motivated/impulsive state (high Hit and Impulsive rates, low Correct Rejection rate and decision bias, and short reaction time), potentially due to being water restricted. As the behavior session progressed, their performance transitioned to a near-optimal regime (high Hit rate, intermediate Correct Rejection rate, high detection sensitivity, and short reaction time). Eventually, mice were much less motivated to perform the task and often disengaged (low Hit and Impulsive rates, high Correct Rejection rate and decision bias, and long reaction time), potentially due to satiety (Fig. 1f). The collective changes of Hit and Correct Rejection rates led to an inverted-U trajectory of overall performance (Fraction Correct, Fig. 1d), which peaked in the middle of a session and declined toward session start and session end. Interestingly, this inverted-U relationship resembles how LC-NE has been hypothesized to modulate task performance (Aston-Jones and Cohen, 2005).

### 3.2 Localized and systemic clonidine treatments similarly impair detection performance

To assess the behavioral effects of suppressing LC activity, we implanted drug delivery cannulae unilaterally in the left LC (contralateral to whisker stimulation) of 6 mice to locally infuse an α2 agonist clonidine (300 nL, 10 mM, 60 nL/min, Fig. 2a). Cannula placement was verified post-hoc to ensure targeted drug administration to the LC (Fig. 2b). Clonidine infusion suppressed LC activity as it reduced c-fos expression in LC neurons (Fig. 2c). On average, c-fos expression was ~40% lower in the clonidine side compared with the contralateral control side (12.7% vs. 19.5%). This reduction was also significant in individual mice (P < 0.01 in 3 out 4 mice, permutation test. Table 1). Saline infusion did not significantly change c-fos expression in the LC (P > 0.05 in 2 mice, permutation test. Table S1). Drug spread was estimated to be ~400 μm (Fig. S3, (St. Peters et al., 2011)). Following clonidine treatments, mice licked less in all trials. As a result, Hit and Impulsive (IS) rates decreased and Correct Rejection (CR) rate increased (Fig. 2d, e). Later in the session, mice showed a tendency of behavioral recovery and re-engaged in the task (Fig. 2d, Fig. S4). Since a typical behavior session in our study lasts 40-50 minutes, this time course is consistent with diminished clonidine effects after ~30 minutes (Abercrombie and Jacobs, 1987; Adams and Foote, 1988; Kalwani et al., 2014). Saline infusion had no effects on behavior (Fig. S5, S6). In addition, in mice where drug infusion was outside of LC we observed minimal behavioral changes (Fig. S7). 5 mM clonidine did not have a significant influence on tactile detection, but the trend is consistent with a dose-dependent effect (Fig. S8). Overall, localized clonidine infusion decreased Hit rate, Impulsive rate and detection sensitivity (d’), elevated Correct Rejection rate, reaction time (RT) and decision bias (c), and impaired task performance (Fig. 2e-g, Fig. S5). Clonidine treated mice behaved as if they were at the end of normal behavior sessions (Fig. 2h). Decreased Impulsive rate, increased reaction time and increased decision bias (changes in c are greater than changes in d’, 1.20 ± 0.15 vs. 0.61 ± 0.10, P = 0.002, n = 10) are all indicative of a motivational shift (Dickinson and Balleine, 1994; Schwarz et al., 2010; Mayrhofer et al., 2013; Berditchevskaia et al., 2016). Thus, we conclude that reduced motivation is the main factor underlying impaired behavior during localized clonidine treatment.

**Figure 2.**
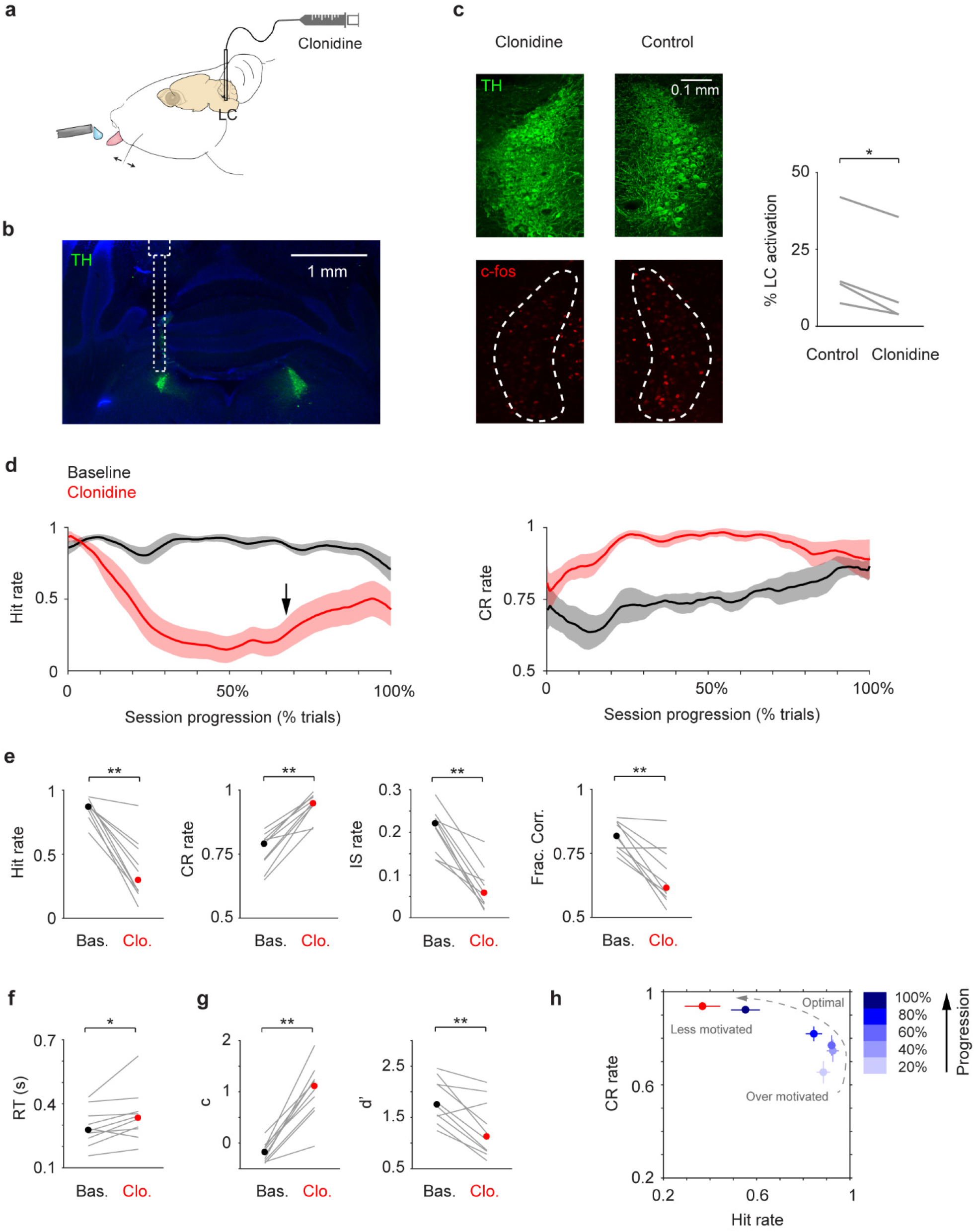
Localized clonidine infusion impairs tactile detection. **a.** Schematic of drug infusion setup. **b.** Histological section showing LC (green) and the tract of the infusion cannula, overlaid with an illustration of cannula placement. **c.** Left: Example c-fos expression (red) in the LC (green) after localized clonidine infusion. The contralateral LC serves as a basal level control. Right: c-fos expression was reduced upon clonidine infusion in 4 awake mice (P = 0.014, two-tailed paired t-test. Cell counts for individual mice are shown in Table 1). % LC activation was defined as the fraction of TH/c-fos double positive cells among TH positive cells. **d.** Mean single-session trajectories for Hit (left) and CR (right) rates during baseline and clonidine sessions (± s.e.m.). Baseline sessions were recorded one day before infusion. Black arrow indicates the onset of Hit rate recovery. **e-g.** Hit rate, CR rate, IS rate, Fraction Correct, RT, decision bias (c) and detection sensitivity (d’) for baseline (black dot, median) and clonidine (red dot, median) sessions. Gray lines indicate individual consecutive two-day, baseline-clonidine pairs. Hit rate, P = 0.002, Signed rank = 55; CR rate, P = 0.002, Signed rank = 0; IS rate, P = 0.002, Signed rank = 55; Frac. Corr., P = 0.0039, Signed rank = 54; RT, P = 0.019, Signed rank = 5; c, P = 0.002, Signed rank = 0; d’, P = 0.0059, Signed rank = 53. n = 10. **h.** CR rate vs. Hit rate trajectory showing clonidine reduces motivation (low Hit rate and high CR rate), which coincides with mouse behavior toward the end of normal baseline sessions. n.s., P > 0.05; * P < 0.05; ** P < 0.01.

**Table 1.** Quantification of c-fos expression to examine the effect of localized clonidine infusion on LC activity in 4 awake mice. Clonidine was infused in the left LC. The right LC serves as a basal level control. Permutation test was performed (10^5^ iterations) to compare c-fos expression levels between the left and right LC in individual mice.

| Clonidine |  |  |  |  |  |  |  |  |
| --- | --- | --- | --- | --- | --- | --- | --- | --- |
| Mouse number | 1 |  | 2 |  | 3 |  | 4 |  |
|  | Left | Right | Left | Right | Left | Right | Left | Right |
| TH positive cells | 424 | 406 | 389 | 252 | 299 | 325 | 169 | 178 |
| TH/c-fos double positive cells | 16 | 56 | 15 | 19 | 106 | 130 | 13 | 26 |
| c-fos expression level (%) | 3.8 | 13.8 | 3.9 | 7.5 | 35.5 | 40.0 | 7.7 | 14.6 |
|  | P < 1e-5 |  | P = 0.0021 |  | P = 0.060 |  | P = 0.0049 |  |

To compare the behavioral effects of localized and systemic drug treatments, we injected clonidine via i.p. (0.05-0.1 mg/kg, (Marzo et al., 2014; Devilbiss, 2019)) in an additional 3 mice. Although systemic drug treatment may affect other areas in the nervous system, the observed behavioral changes resembled localized infusion (reduced Hit rate, Impulsive rate and detection sensitivity, elevated Correct Rejection rate, reaction time and decision bias, Fig. 3a-c). Saline injection did not affect behavior (Fig. S9).

**Figure 3.**
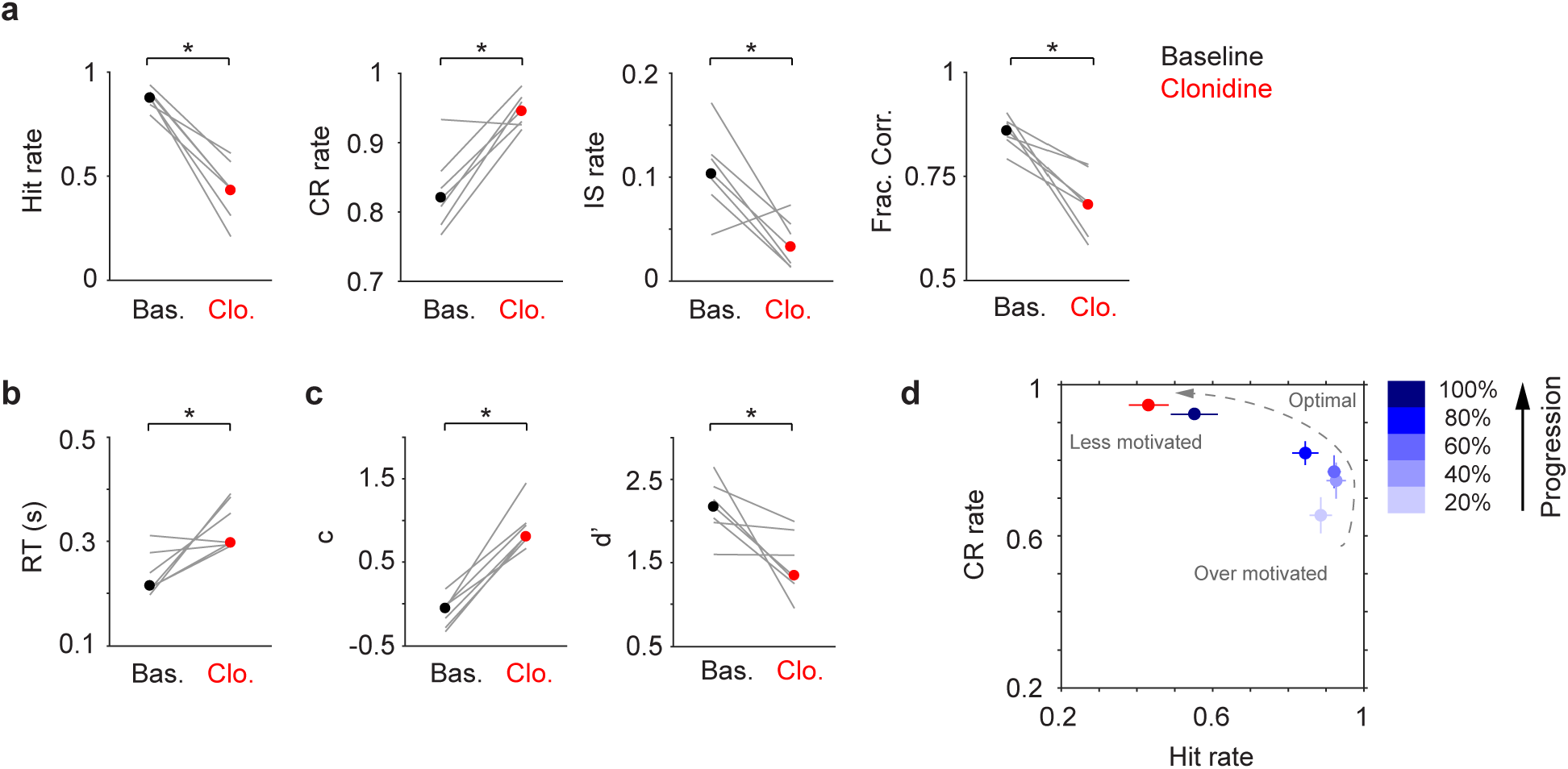
Systemic clonidine treatment impairs tactile detection. **a-c.** Hit rate, CR rate, IS rate, Fraction Correct, RT, decision bias (c) and detection sensitivity (d’) for baseline (black dot, median) and clonidine (red dot, median) sessions. Gray lines indicate individual consecutive two-day, baseline-clonidine pairs. Hit rate, P = 0.016, Signed rank = 28; CR rate, P = 0.031, Signed rank = 1; IS rate, P = 0.031, Signed rank = 27; Frac. Corr., P = 0.016, Signed rank = 28; RT, P = 0.031, Signed rank = 1; c, P = 0.016, Signed rank = 0; d’, P = 0.016, Signed rank = 28. n = 7. **d.** CR rate vs. Hit rate trajectory showing clonidine reduces motivation (low Hit rate and high CR rate), similar to localized infusion in **Fig. 2h**. * P < 0.05.

To conclude, we found that both localized and systemic clonidine treatments (decreasing LC activity) impaired task performance in a similar fashion, i.e., by reducing motivation (Fig. 2h, Fig. 3d).

### 3.3 Localized and systemic yohimbine treatments differently affect detection performance

Next, to assess the behavioral effects of enhancing LC activity, we locally infused an α2 antagonist yohimbine (300 nL, 10 mM, 60 nL/min) in the left LC of 7 mice. Localized yohimbine administration enhanced LC activity as it increased c-fos expression in LC neurons (Fig. 4a). On average, c-fos expression was ~100% higher in the yohimbine side compared with the contralateral control side (38.6% vs. 19.9%). This effect was significant in individual mice (P < 1e-5 in all 5 mice, permutation test. Table 2). Interestingly, we did not observe any changes in Hit rate after yohimbine infusion, but Correct Rejection (CR) rate was significantly increased, accompanied with a reduction of Impulsive (IS) rate (Fig. 4b, c, Fig. S10). We note that later in the session Correct Rejection rate returned to baseline levels (after ~30 minutes, Fig. 4b), consistent with the time course of diminished yohimbine effects (Andén et al., 1982). However, it has also been reported that elevated LC baseline firing could be sustained up to 60 minutes upon yohimbine administration (Rasmussen and Jacobs, 1986). Saline treatment did not affect behavior (Fig. S10, S11). 20 mM yohimbine had a similar influence on behavior as 10 mM, and the trend is consistent with a dose-dependent effect (Fig. S12). However, 20 mM yohimbine appeared to induce transient behavioral arrests during the initial 50-100 trials (data not shown), implying that this dose over-activates LC (Carter et al., 2010). Overall, the primary behavioral effect of localized yohimbine treatment was an improvement of task performance as mice could better withhold licking in NoGo trials and were less prone to False Alarms (Fig. 4b-d, Fig. S10), resembling their peak performance in the middle of normal behavior sessions (Fig. 4f). Yohimbine did not affect decision bias but significantly increased detection sensitivity (Fig. 4e, Fig. S10), which suggests that the behavioral improvement is not simply a result of an overall increase of arousal (which would be reflected by significant decreases in decision bias (Gelbard-Sagiv et al., 2018)), but more specifically of enhanced sensory processing (e.g., increased signal-to-noise ratio).

**Table 2.** Quantification of c-fos expression to examine the effect of localized yohimbine infusion on LC activity in 5 mice. Yohimbine was infused in the left LC (Mouse 1 and 2: anesthesia; Mouse 3-5: awake). The right LC serves as a basal level control. Permutation test was performed (10^5^ iterations) to compare c-fos expression levels between the left and right LC in individual mice.

| Yohimbine |  |  |  |  |  |  |  |  |  |  |
| --- | --- | --- | --- | --- | --- | --- | --- | --- | --- | --- |
| Mouse number | 1 |  | 2 |  | 3 |  | 4 |  | 5 |  |
|  | Left | Right | Left | Right | Left | Right | Left | Right | Left | Right |
| TH positive cells | 531 | 536 | 416 | 402 | 100 | 196 | 355 | 382 | 501 | 458 |
| TH/c-fos double positive cells | 209 | 106 | 112 | 12 | 52 | 51 | 191 | 153 | 104 | 48 |
| c-fos expression level (%) | 39.4 | 19.8 | 26.9 | 3.0 | 52.0 | 26.0 | 53.8 | 40.1 | 20.8 | 10.5 |
|  | P < 1e-5 |  | P < 1e-5 |  | P < 1e-5 |  | P < 1e-5 |  | P < 1e-5 |  |

**Figure 4.**
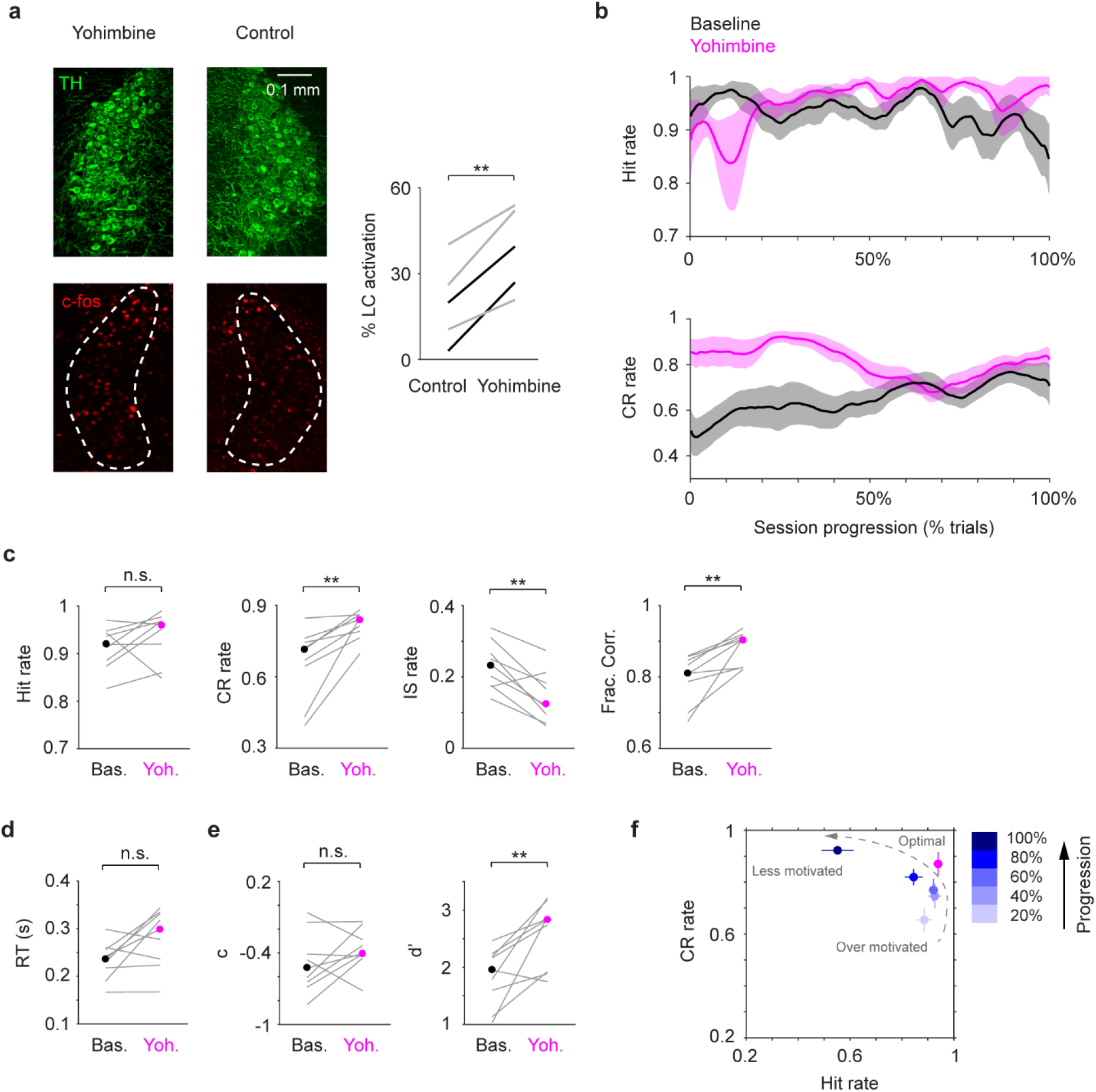
Localized yohimbine infusion improves tactile detection. **a.** Left: Example c-fos expression (red) in the LC (green) after localized yohimbine infusion. The contralateral LC serves as a basal level control. Right: c-fos expression was enhanced upon yohimbine infusion in 5 mice (P = 0.0033, two-tailed paired t-test. 2 under anesthesia, black lines; 3 during wakefulness, gray lines. Cell counts for individual mice are shown in Table 2). % LC activation was defined as the fraction of TH/c-fos double positive cells among TH positive cells. **b.** Mean single-session trajectories for Hit (top) and CR (bottom) rates during baseline and yohimbine sessions (± s.e.m.). Baseline sessions were recorded one day before infusion. **c-e.** Hit rate, CR rate, IS rate, Fraction Correct, RT, decision bias (c) and detection sensitivity (d’) for baseline (black dot, median) and clonidine (magenta dot, median) sessions. Gray lines indicate individual consecutive two-day, baseline-yohimbine pairs. Hit rate, P = 0.20, Signed rank = 11; CR rate, P = 0.0039, Signed rank = 0; IS rate, P = 0.0078, Signed rank = 44; Frac. Corr., P = 0.0039, Signed rank = 0; RT, P = 0.074, Signed rank = 7; c, P = 0.30, Signed rank = 11; d’, P = 0.0078, Signed rank = 1. n = 9. **f.** CR rate vs. Hit rate trajectory showing yohimbine transitioned mouse behavior to a near-optimal regime (high Hit rate and high CR rate), similar to mouse behavior around the middle of normal baseline sessions. n.s., P > 0.05; ** P < 0.01.

To compare the behavioral effects of localized and systemic drug treatments, we injected yohimbine via i.p. in 5 mice (2 mg/kg, (Rasmussen and Jacobs, 1986)). Contrary to localized infusion, systemically treated mice were less capable of withholding licks during the waiting periods as well as in NoGo trials, resulting in increased Impulsive rate and reduced Correct Rejection rate, decision bias and detection sensitivity (Fig. 5a-c). These behavioral changes are consistent with an increase of impulsivity, and mice behaved as if they were at the beginning of normal behavior sessions (Fig. 5d).

**Figure 5.**
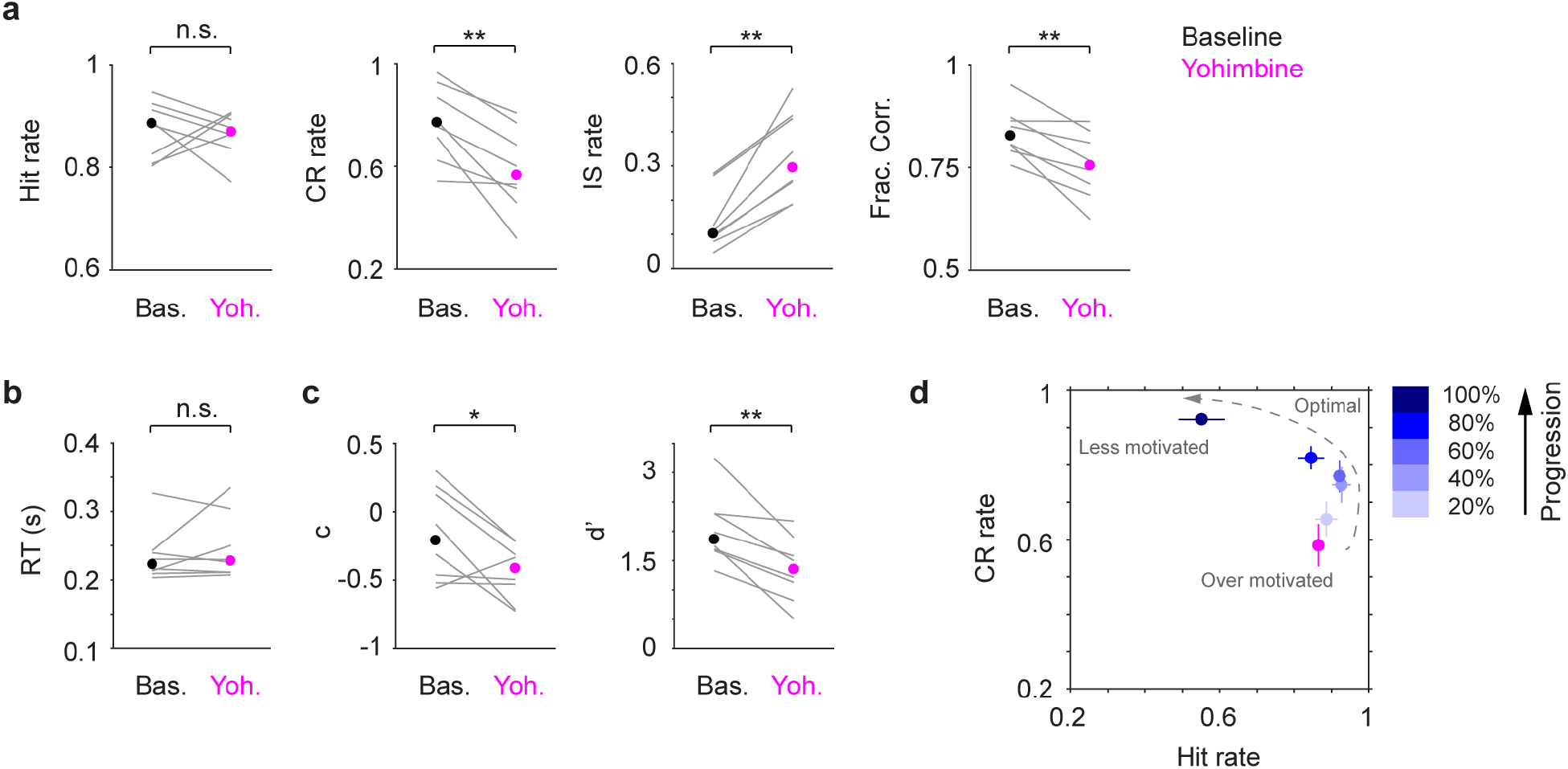
Systemic yohimbine treatment impairs tactile detection. **a-c.** Hit rate, CR rate, IS rate, Fraction Correct, RT, decision bias (c) and detection sensitivity (d’) for baseline (black dot, mean) and yohimbine (magenta dot, mean) sessions. Gray lines indicate individual consecutive two-day, baseline-yohimbine pairs. Hit rate, P = 1, Signed rank = 18; CR rate, P = 0.0078, Signed rank = 36; IS rate, P = 0.0078, Signed rank = 0; Frac. Corr., P = 0.0078, Signed rank = 36; RT, P = 0.84, Signed rank = 16; c, P = 0.039, Signed rank = 33; d’, P = 0.0078, Signed rank = 36. n = 8. **d.** CR rate vs. Hit rate trajectory showing yohimbine promotes impulsivity (high Hit rate and low CR rate), which coincides with mouse behavior at the beginning of normal baseline sessions. n.s., P > 0.05; * P < 0.05; ** P < 0.01.

To conclude, we found that localized and systemic yohimbine treatments (increasing LC activity) exerted opposing behavioral effects. Localized infusion improved tactile detection, and mice achieved near-optimal performance (Fig. 4f). In contrast, systemic treatment impaired performance by promoting impulsivity (Fig. 5d).

## 4. Discussion

The current study is one of the first to investigate how bidirectional perturbations of LC-NE affect quantitative perceptual task performance. We found that localized and systemic pharmacological suppression of LC-NE similarly impaired tactile detection (decreased Hit and Impulsive rates, elevated Correct Rejection rate and decision bias, and prolonged reaction time), suggesting that a major site of action during systemic clonidine treatment is the LC.

Our results support previous findings that suppressing LC-NE signaling decreases arousal, promotes sleep, and slows down reaction time (Sarro et al., 1987; Halliday et al., 1989; Berridge et al., 1993; Turetsky and Fein, 2002; Hou et al., 2005; Carter et al., 2010). Given that the main effect of suppressing LC-NE is to reduce arousal/motivation, the behavioral impairment is likely to be task-independent. A recent study showed that systemic clonidine did not affect decision bias (Gelbard-Sagiv et al., 2018). In this study human subjects were instructed to adjust their preparedness before initiating a new trial, which possibly engaged other arousal-promoting circuits (e.g., the cholinergic system (McGaughy et al., 1996)) to compensate the clonidine-induced decline of arousal/motivation (Thiele and Bellgrove, 2018).

In terms of activation, we found that localized yohimbine infusion in the LC improved tactile detection (increased Correct Rejection rate and detection sensitivity, and reduced Impulsive rate), while systemic yohimbine treatment impaired behavior (elevated Impulsive rate, and decreased Correct Rejection rate, decision bias and detection sensitivity). Our findings are consistent with others showing that systemic yohimbine increased impulsivity (e.g., (Swann et al., 2005, 2013; Sun et al., 2010)). The different behavioral effects between localized and systemic treatments suggest that increased impulsivity is likely due to yohimbine acting on presynaptic and postsynaptic α2 ARs (Starke et al., 1975; Szemeredi et al., 1991; Arnsten and Cai, 1993) in “off-target” α2-expressing regions, such as noradrenergic neurons in the nucleus of the solitary tract (Van Bockstaele et al., 1999; Kirouac, 2015), and the prefrontal cortex (Solanto, 1998; Arnsten, 2000; Ramos and Arnsten, 2007; Sun et al., 2010; Janitzky et al., 2015). It should be noted that yohimbine also has pronounced affinity to 5-HT1 receptors and dopamine D2 receptors (Millan et al., 2000). In addition, activating LC via localized or systemic administration of corticotropin-releasing factors differently affected rats performing an attention set shifting task (Snyder et al., 2012). Together, these findings strongly suggest that systemic yohimbine treatment, or in general non-specific NE activation, cannot be interpreted as specific manipulation of the LC-NE circuit.

Importantly, whether systemic (non-specific) NE activation impairs or improves task performance likely depends on the brain regions, the receptors (adrenergic and non-adrenergic), and the type of behavior task involved. For example, during systemic administration of the psychostimulant methylphenidate (MPH, an NE-DA reuptake inhibitor), enhanced NE release acting on α1 ARs in the prefrontal cortex was reasoned to underlie the dose-dependent changes in rats performing a sustained attention task (Berridge et al., 2006, 2012; Andrzejewski et al., 2014; Spencer et al., 2015; Berridge and Spencer, 2016). On the other hand, activation of the prefrontal α2 ARs and dopamine D1 receptors during MPH administration contributed to the improved performance in a spatial working memory task (e.g., (Arnsten and Dudley, 2005; Berridge et al., 2006)).

Our study could have implications for several neurological disorders, including attention-deficit hyperactivity disorder (ADHD), for which one of the major diagnostic criteria is impulsive behavior (Castellanos and Tannock, 2002). In children performing a Go/NoGo learning task, those diagnosed with ADHD had a higher False Alarm rate than controls (e.g., (Iaboni et al., 1995)). Mice with ADHD-phenotypes also exhibited higher False Alarm and Impulsive rates during Go/NoGo motor tests (Majdak et al., 2016). Interestingly, this impulsive/distractible response has been linked to high tonic LC activity (Rajkowski et al., 1994; Usher et al., 1999; Aston-Jones and Cohen, 2005). Consistent with these findings, clonidine, and possibly other α2 agonists, can suppress LC activity and reduce impulsivity (Mangeot et al., 2001; Berridge and Waterhouse, 2003).

We found that unilateral LC perturbation (contralateral to whisker stimulation) is sufficient to produce pronounced behavioral changes. Since unilateral LC suppression mainly reduced arousal/motivation, it suggests that this manipulation affects arousal-related circuits downstream of LC, such as the basal forebrain cholinergic system and the preoptic area of the hypothalamus (Jones and Moore, 1977; España and Berridge, 2006). Thus, we anticipate that the behavioral impairment is laterality-independent, i.e., suppressing the LC ipsilateral to whisker stimulation would similarly reduce arousal/motivation. We found that unilateral LC activation improves tactile detection. In our behavior task, the right C2 whisker was stimulated, and yohimbine was infused in the left LC. In rodents, the ascending whisker information is fully crossed in somatosensory thalamus and cortex (Diamond et al., 2008), which in turn receive extensive innervations from the ipsilateral LC (Simpson et al., 1997). Since unilateral LC activation improves detection sensitivity (d’) while leaving decision bias (c) unaffected, our results imply that activating the left LC enhances the representation of the contralateral (right) whisker stimulation to improve task performance, potentially through NE modulating the ipsilateral (left) somatosensory thalamus and/or somatosensory cortex. This interpretation is in line with previous results showing that enhanced LC-NE signaling improves sensory processing in somatosensation-related areas (e.g., (Lecas, 2004; Devilbiss et al., 2006; Hirata et al., 2006; Vazey et al., 2018)). We anticipate that stimulating the right LC (ipsilateral to whisker stimulation) would not produce similar behavioral effects, and that the behavioral improvement is laterality- and task-dependent (e.g., perceptual vs. non-perceptual). However, it remains a possibility that unilateral LC activation could enhance bilateral LC responses (Marzo et al., 2014), and stimulating the right LC could produce similar behavioral improvement. Future experiments are needed to test these hypotheses.

Our localized yohimbine results support two recent studies testing how activating LC-NE affects perceptual task performance. In one, LC was optogenetically activated in rats performing a tactile frequency discrimination task (Rodenkirch et al., 2019). In another, LC-NE signaling was enhanced by using a selective NE reuptake inhibitor in human subjects performing visual detection/discrimination tasks (Gelbard-Sagiv et al., 2018). Regardless of the differences in species and perturbation methods, activating LC-NE improves sensitivity (d’) and performance, suggesting that the behavioral enhancement is more specific to LC-NE acting on sensory processing-related areas. Future work is needed to examine how LC projections in different somatosensory areas differentially contribute to tactile perception, and how perturbing LC-NE modulates other types of behavioral tasks.

## Acknowledgements

We thank B. A. Bari, M. Riccomagno, P. Hickmott, V. Santhakumar and members of the Yang lab for comments on the manuscript, and L. Graham for instrument fabrication. This work was supported by UCR startup (S.H.Y, H.Y.), UC Regents’ Faculty Fellowship (H.Y.), Klingenstein-Simons Fellowship Awards in Neuroscience (H.Y.), and National Institute of Neurological Disorders and Stroke 1R01NS107355 and 1R01NS112200 (H.Y.).

## Author contributions

J.M.L., Y.S. and H.Y. planned the project. J.M.L., Y.S., and L.S.T. performed experiments. Q.A.N. and S.H.Y. helped with c-fos staining and quantification. J.M.L. and H.Y. built apparatus, analyzed data and wrote the manuscript with comments from all authors.

## References

Abercrombie ED, Jacobs BL (1987) Microinjected clonidine inhibits noradrenergic neurons of the locus coeruleus in freely moving cats. Neurosci Lett 76:203–208.

Acheson A, Zigmond M, Stricker E (1980) Compensatory increase in tyrosine hydroxylase activity in rat brain after intraventricular injections of 6-hydroxydopamine. Science (80-) 207:537–540.

Adams LM, Foote SL (1988) Effects of locally infused pharmacological agents on spontaneous and sensory-evoked activity of locus coeruleus neurons. Brain Res Bull 21:395–400.

Aghajanian GK, VanderMaelen CP (1982) alpha 2-adrenoceptor-mediated hyperpolarization of locus coeruleus neurons: intracellular studies in vivo. Science 215:1394–1396.

Aghajanian GK, Wang YY (1987) Common α2- and opiate effector mechanisms in the locus coeruleus: intracellular studies in brain slices. Neuropharmacology 26:793–799.

Andén NE, Pauksens K, Svensson K (1982) Selective blockade of brain α2-autoreceptors by yohimbine: Effects on motor activity and on turnover of noradrenaline and dopamine. J Neural Transm 55:111–120.

Andrzejewski ME, Spencer RC, Harris RL, Feit EC, McKee BL, Berridge CW (2014) The effects of clinically relevant doses of amphetamine and methylphenidate on signal detection and DRL in rats. Neuropharmacology 79:634–641.

Arnsten AFT (2000) Through the Looking Glass: Differential Noradenergic Modulation of Prefrontal Cortical Function. Neural Plast 7:133–146.

Arnsten AFT, Cai JX (1993) Postsynaptic alpha-2 receptor stimulation improves memory in aged monkeys: Indirect effects of yohimbine versus direct effects of clonidine. Neurobiol Aging 14:597–603.

Arnsten AFT, Dudley AG (2005) Methylphenidate improves prefrontal cortical cognitive function through α2 adrenoceptor and dopamine D1 receptor actions: Relevance to therapeutic effects in Attention Deficit Hyperactivity Disorder. Behav Brain Funct 1:1–9.

Aston-Jones G, Cohen JD (2005) An integrative theory of locus coeruleus-norepinephrine function: adaptive gain and optimal performance. Annu Rev Neurosci 28:403–450.

Bankhead P, Loughrey MB, Fernández JA, Dombrowski Y, McArt DG, Dunne PD, McQuaid S, Gray RT, Murray LJ, Coleman HG, James JA, Salto-Tellez M, Hamilton PW (2017) QuPath: Open source software for digital pathology image analysis. Sci Rep 7:1–7.

Berditchevskaia A, Cazé RD, Schultz SR (2016) Performance in a GO/NOGO perceptual task reflects a balance between impulsive and instrumental components of behaviour. Sci Rep 6:1–15.

Berridge CW, Devilbiss DM, Andrzejewski ME, Arnsten AFT, Kelley AE, Schmeichel B, Hamilton C, Spencer RC (2006) Methylphenidate Preferentially Increases Catecholamine Neurotransmission within the Prefrontal Cortex at Low Doses that Enhance Cognitive Function. Biol Psychiatry 60:1111–1120.

Berridge CW, Page ME, Valentino RJ, Foote SL (1993) Effects of locus coeruleus inactivation on electroencephalographic activity in neocortex and hippocampus. Neuroscience 55:381–393.

Berridge CW, Shumsky JS, Andrzejewski ME, McGaughy JA, Spencer RC, Devilbiss DM, Waterhouse BD (2012) Differential sensitivity to psychostimulants across prefrontal cognitive tasks: Differential involvement of noradrenergic ?? 1- and ?? 2-receptors. Biol Psychiatry 71:467–473.

Berridge CW, Spencer RC (2016) Differential cognitive actions of norepinephrine a2 and a1 receptor signaling in the prefrontal cortex. Brain Res 1641:189–196.

Berridge CW, Waterhouse BD (2003) The locus coeruleus - noradrenergic system: modulation of behavioral state and state-dependent cognitive processes. Brain Res Rev 42:33–84.

Carter ME, Yizhar O, Chikahisa S, Nguyen H, Adamantidis A, Nishino S, Deisseroth K, de Lecea L (2010) Tuning arousal with optogenetic modulation of locus coeruleus neurons. Nat Neurosci 13:1526–1533.

Castellanos FX, Tannock R (2002) Neuroscience of attention-deficit/hyperactivity disorder: The search for endophenotypes. Nat Rev Neurosci 3:617–628.

Cedarbaum JM, Aghajanian GK (1976) Noradrenergic neurons of the locus coeruleus: inhibition by epinephrine and activation by the α-antagonist piperoxane. Brain Res 112:413–419.

Cedarbaum JM, Aghajanian GK (1977) Catecholamine receptors on locus coeruleus neurons: Pharmacological characterization. Eur J Pharmacol 44:375–385.

Devilbiss DM (2019) Consequences of tuning network function by tonic and phasic locus coeruleus output and stress: Regulating detection and discrimination of peripheral stimuli. Brain Res 1709:16–27.

Devilbiss DM, Page ME, Waterhouse BD (2006) Locus ceruleus regulates sensory encoding by neurons and networks in waking animals. J Neurosci 26:9860–9872.

Devilbiss DM, Waterhouse BD (2004) The effects of tonic locus ceruleus output on sensory-evoked responses of ventral posterior medial thalamic and barrel field cortical neurons in the awake rat. J Neurosci 24:10773–10785.

Diamond ME, Von Heimendahl M, Knutsen PM, Kleinfeld D, Ahissar E (2008) “Where” and “what” in the whisker sensorimotor system. Nat Rev Neurosci 9:601–612.

Dickinson A, Balleine B (1994) Motivational control of goal-directed action. Anim Learn Behav 22:1–18.

Doucette W, Milder J, Restrepo D (2007) Adrenergic modulation of olfactory bulb circuitry affects odor discrimination. Learn & Mem 14:539–547.

Escanilla O, Arrellanos A, Karnow A, Ennis M, Linster C (2010) Noradrenergic modulation of behavioral odor detection and discrimination thresholds in the olfactory bulb. Eur J Neurosci 32:458–468.

España RA, Berridge CW (2006) Organization of noradrenergic efferents to arousal-related basal forebrain structures. J Comp Neurol 496:668–683.

Foote SL, Freedman R, Oliver AP (1975) Effects of putative neurotransmitters on neuronal activity in monkey auditory cortex. Brain Res 86:229–242.

Gelbard-Sagiv H, Magidov E, Sharon H, Hendler T, Nir Y (2018) Noradrenaline Modulates Visual Perception and Late Visually Evoked Activity. Curr Biol 28:2239–2249.e6.

Green DM, Swets JA (1966) Signal detection theory and psychophysics. John Wiley and Sons Inc.

Guo Z V., Hires SA, Li N, O’Connor DH, Komiyama T, Ophir E, Huber D, Bonardi C, Morandell K, Gutnisky D, Peron S, Xu N, Cox J, Svoboda K (2014) Procedures for Behavioral Experiments in Head-Fixed Mice Simon SA, ed. PLoS One 9:e88678.

Halliday R, Callaway E, Lannon R (1989) The effects of clonidine and yohimbine on human information processing. Psychopharmacology (Berl) 99:563–566.

Harik S, Duckrow R, LaManna J, Rosenthal M, Sharma V, Banerjee S (1981) Cerebral compensation for chronic noradrenergic denervation induced by locus ceruleus lesion: recovery of receptor binding, isoproterenol-induced adenylate cyclase activity, and oxidative metabolism. J Neurosci 1:641–649.

Herr NR, Park J, McElligott Z a., Belle a. M, Carelli RM, Wightman RM (2012) In vivo voltammetry monitoring of electrically evoked extracellular norepinephrine in subregions of the bed nucleus of the stria terminalis. J Neurophysiol 107:1731–1737.

Hirata A, Aguilar J, Castro-Alamancos MA (2006) Noradrenergic activation amplifies bottom-up and top-down signal-to-noise ratios in sensory thalamus. J Neurosci 26:4426–4436.

Hou RH, Freeman C, Langley RW, Szabadi E, Bradshaw CM (2005) Does modafinil activate the locus coeruleus in man? Comparison of modafinil and clonidine on arousal and autonomic functions in human volunteers. Psychopharmacology (Berl) 181:537–549.

Iaboni F, Douglas VI, Baker AG (1995) Effects of reward and response costs on inhibition in ADHD children. J Abnorm Psychol 104:232–240.

Janitzky K, Lippert MT, Engelhorn A, Tegtmeier J, Goldschmidt J, Heinze H-J, Ohl FW (2015) Optogenetic silencing of locus coeruleus activity in mice impairs cognitive flexibility in an attentional set-shifting task. Front Behav Neurosci 9:1–8.

Jones BE, Moore RY (1977) Ascending projections of the locus coeruleus in the rat. II. Autoradiographic study. Brain Res 127:23–53.

Kalwani RM, Joshi S, Gold JI (2014) Phasic Activation of Individual Neurons in the Locus Ceruleus/Subceruleus Complex of Monkeys Reflects Rewarded Decisions to Go But Not Stop. J Neurosci 34:13656–13669.

Kasamatsu T, Heggelund P (1982) Single cell responses in cat visual cortex to visual stimulation during iontophoresis of noradrenaline. Exp brain Res 45:317–327.

Kirouac GJ (2015) Placing the paraventricular nucleus of the thalamus within the brain circuits that control behavior. Neurosci Biobehav Rev 56:315–329.

Lecas JC (2004) Locus coeruleus activation shortens synaptic drive while decreasing spike latency and jitter in sensorimotor cortex. Implications for neuronal integration. Eur J Neurosci 19:2519–2530.

Lee SH, Dan Y (2012) Neuromodulation of Brain States. Neuron 76:109–222.

Majdak P, Ossyra JR, Ossyra JM, Cobert AJ, Hofmann GC, Tse S, Panozzo B, Grogan EL, Sorokina A, Rhodes JS (2016) A new mouse model of ADHD for medication development. Sci Rep 6:1–18.

Manella LC, Petersen N, Linster C (2017) Stimulation of the Locus Ceruleus Modulates Signal-to-Noise Ratio in the Olfactory Bulb. J Neurosci 37:11605–11615.

Mangeot SD, Miller LJ, McIntosh DN, McGrath-Clarke J, Simon J, Hagerman RJ, Goldson E (2001) Sensory modulation dysfunction in children with attention-deficit-hyperactivity disorder. Dev Med Child Neurol 43:399.

Martins ARO, Froemke RC (2015) Coordinated forms of noradrenergic plasticity in the locus coeruleus and primary auditory cortex. Nat Neurosci 18:1–12.

Marzo A, Totah NK, Neves RM, Logothetis NK, Eschenko O (2014) Unilateral electrical stimulation of rat locus coeruleus elicits bilateral response of norepinephrine neurons and sustained activation of medial prefrontal cortex. J Neurophysiol 111:2570–2588.

Mayrhofer JM, Skreb V, von der Behrens W, Musall S, Weber B, Haiss F (2013) Novel two-alternative forced choice paradigm for bilateral vibrotactile whisker frequency discrimination in head-fixed mice and rats. J Neurophysiol 109:273–284.

McBurney-Lin J, Lu J, Zuo Y, Yang H (2019) Locus coeruleus-norepinephrine modulation of sensory processing and perception: A focused review. Neurosci Biobehav Rev 105:190–199.

McCune SK, Voigt MM, Hill‡ JM (1993) Expression of multiple alpha adrenergic receptor subtype messenger RNAs in the adult rat brain. Neuroscience 57:143–151.

McGaughy J, Kaiser T, Sarter M (1996) Behavioral vigilance following infusions of 192 IgG-saporin into the basal forebrain: Selectivity of the behavioral impairment and relation to cortical AChE-positive fiber density. Behav Neurosci 110:247–265.

Millan MJ, Newman-Tancredi A, Audinot V, Cussac D, Lejeune F, Nicolas JP, Cogé F, Galizzi JP, Boutin JA, Rivet JM, Dekeyne A, Gobert A (2000) Agonist and antagonist actions of yohimbine as compared to fluparoxan at α2-adrenergic receptors (AR)s, serotonin (5-HT)(1A), 5-HT(1B), 5-HT(1D) and dopamine D2 and D3 receptors. Significance for the modulation of frontocortical monoaminergic transmission. Synapse 35:79–95.

Morrison JH, Foote SL (1986) Noradrenergic and serotoninergic innervation of cortical, thalamic, and tectal visual structures in old and new world monkeys. J Comp Neurol 243:117–138.

Navarra RL, Clark BD, Gargiulo AT, Waterhouse BD (2017) Methylphenidate Enhances Early-Stage Sensory Processing and Rodent Performance of a Visual Signal Detection Task. Neuropsychopharmacology 42:1326–1337.

Neves RM, van Keulen S, Yang M, Logothetis NK, Eschenko O (2018) Locus coeruleus phasic discharge is essential for stimulus-induced gamma oscillations in the prefrontal cortex. J Neurophysiol 119:904–920.

Nicholas AP, Pieribone V, Hokfelt T (1993) Distributions of mRNAs for alpha-2 adrenergic receptor subtypes in rat brain: an in situ hybridization study. J Comp Neurol 328:575–594.

Raiteri M, Maura G, Versace P (1983) Functional evidence for two stereochemically different alpha-2 adrenoceptors regulating central norepinephrine and serotonin release. J Pharmacol Exp Ther 224:679–684.

Rajkowski J, Kubiak P, Aston-Jones G (1994) Locus coeruleus activity in monkey: Phasic and tonic changes are associated with altered vigilance. Brain Res Bull 35:607–616.

Ramos BP, Arnsten AFT (2007) Adrenergic pharmacology and cognition: Focus on the prefrontal cortex. Pharmacol Ther 113:523–536.

Rasmussen K, Jacobs BL (1986) Single unit activity of locus coeruleus neurons in the freely moving cat. Brain Res 371:335–344.

Rho HJ, Kim JH, Lee SH (2018) Function of selective neuromodulatory projections in the mammalian cerebral cortex: Comparison between cholinergic and noradrenergic systems. Front Neural Circuits 12:1–13.

Robertson SD, Plummer NW, de Marchena J, Jensen P (2013) Developmental origins of central norepinephrine neuron diversity. Nat Neurosci 16:1016–1023.

Rodenkirch C, Liu Y, Schriver BJ, Wang Q (2019) Locus coeruleus activation enhances thalamic feature selectivity via norepinephrine regulation of intrathalamic circuit dynamics. Nat Neurosci 22:120–133.

Sara SJ (2009) The locus coeruleus and noradrenergic modulation of cognition. Nat Rev Neurosci 10:211–223.

Sara SJ, Bouret S (2012) Orienting and Reorienting: The Locus Coeruleus Mediates Cognition through Arousal. Neuron 76:130–141.

Sarro GB, Ascioti C, Froio F, Libri V, Nisticò G (1987) Evidence that locus coeruleus is the site where clonidine and drugs acting at α1- and α2-adrenoceptors affect sleep and arousal mechanisms. Br J Pharmacol 90:675–685.

Schindelin J, Arganda-Carreras I, Frise E, Kaynig V, Longair M, Pietzsch T, Preibisch S, Rueden C, Saalfeld S, Schmid B, Tinevez JY, White DJ, Hartenstein V, Eliceiri K, Tomancak P, Cardona A (2012) Fiji: An open-source platform for biological-image analysis. Nat Methods 9:676–682.

Schwarz C, Hentschke H, Butovas S, Haiss F, Stüttgen MC, Gerdjikov T V., Bergner CG, Waiblinger C (2010) The head-fixed behaving rat—Procedures and pitfalls. Somatosens Mot Res 27:131–148.

Simpson KL, Altman DW, Wang L, Kirifides ML, Lin RCS, Waterhouse BD (1997) Lateralization and functional organization of the locus coeruleus projection to the trigeminal somatosensory pathway in rat. J Comp Neurol 385:135–147.

Simson PE, Weiss JM (1987) Alpha-2 receptor blockade increases responsiveness of locus coeruleus neurons to excitatory stimulation. J Neurosci 7:1732–1740.

Snyder K, Wang WW, Han R, McFadden K, Valentino RJ (2012) Corticotropin-releasing factor in the norepinephrine nucleus, locus coeruleus, facilitates behavioral flexibility. Neuropsychopharmacology 37:520–530.

Solanto M V. (1998) Neuropsychopharmacological mechanisms of stimulant drug action in attention-deficit hyperactivity disorder: A review and integration. Behav Brain Res 94:127–152.

Spencer RC, Devilbiss DM, Berridge CW (2015) The cognition-enhancing effects of psychostimulants involve direct action in the prefrontal cortex. Biol Psychiatry 77:940–950.

St. Peters M, Demeter E, Lustig C, Bruno JP, Sarter M (2011) Enhanced Control of Attention by Stimulating Mesolimbic-Corticopetal Cholinergic Circuitry. J Neurosci 31:9760–9771.

Starke K, Borowski E, Endo T (1975) Preferential blockade of presynaptic α-adrenoceptors by yohimbine. Eur J Pharmacol 34:385–388.

Sun HS, Green TA, Theobald DEH, Birnbaum SG, Graham DL, Zeeb FD, Nestler EJ, Winstanley CA (2010) Yohimbine Increases Impulsivity Through Activation of cAMP Response Element Binding in the Orbitofrontal Cortex. Biol Psychiatry 67:649–656.

Swann AC, Birnbaum D, Jagar AA, Dougherty DM, Moeller FG (2005) Acute yohimbine increases laboratory-measured impulsivity in normal subjects. Biol Psychiatry 57:1209–1211.

Swann AC, Lijffijt M, Lane SD, Cox B, Steinberg JL, Moeller FG (2013) Norepinephrine and impulsivity: Effects of acute yohimbine. Psychopharmacology (Berl) 229:83–94.

Szemeredi K, Komoly S, Kopin IJ, Bagdy G, Keiser HR, Goldstein DS (1991) Simultaneous measurement of plasma and brain extracellular fluid concentrations of catechols after yohimbine administration in rats. Brain Res 542:8–14.

Thiele A, Bellgrove MA (2018) Neuromodulation of Attention. Neuron 97:769–785.

Turetsky BI, Fein G (2002) α2-noradrenergic effects on ERP and behavioral indices of auditory information processing. Psychophysiology 39:147–157.

Usher M, Cohen JD, Servan-Schreiber D, Rajkowski J, Aston-Jones G (1999) The role of locus coeruleus in the regulation of cognitive performance. Science 283:549–554.

Valentini V, Frau R, Di Chiara G (2004) Noradrenaline transporter blockers raise extracellular dopamine in medial prefrontal but not parietal and occipital cortex: Differences with mianserin and clozapine. J Neurochem 88:917–927.

Van Bockstaele EJ, Peoples J, Telegan P (1999) Efferent projections of the nucleus of the solitary tract to peri-Locus coeruleus dendrites in rat brain: Evidence for a monosynaptic pathway. J Comp Neurol 412:410–428.

Vazey EM, Moorman DE, Aston-Jones G (2018) Phasic locus coeruleus activity regulates cortical encoding of salience information. Proc Natl Acad Sci 115:E9439–E9448.

Waterhouse BD, Moises HC, Woodward DJ (1980) Noradrenergic modulation of somatosensory cortical neuronal responses to lontophoretically applied putative neurotransmitters. Exp Neurol 69:30–49.

Waterhouse BD, Navarra RL (2019) The locus coeruleus-norepinephrine system and sensory signal processing: A historical review and current perspectives. Brain Res 1709:1–15.

Yang H, Kwon SE, Severson KS, O’Connor DH (2016) Origins of choice-related activity in mouse somatosensory cortex. Nat Neurosci 19:127–134.

