## Supplemental information for "Bidirectional pharmacological perturbations of the noradrenergic system differentially affect tactile detection"

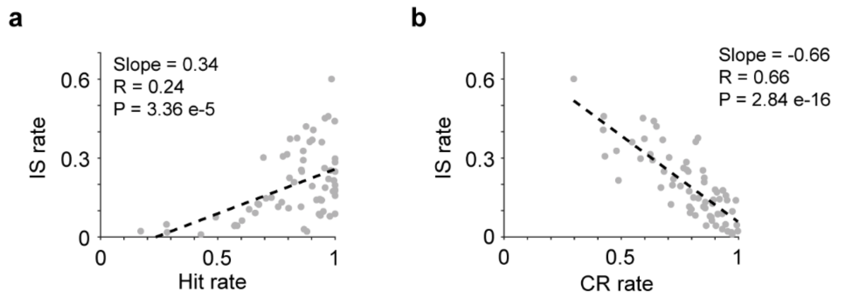


**Figure S1.** Impulsive (IS) rate was positively correlated with Hit rate (**a**), and negatively correlated with Correct Rejection (CR) rate (**b**). Dashed lines: linear fitting.

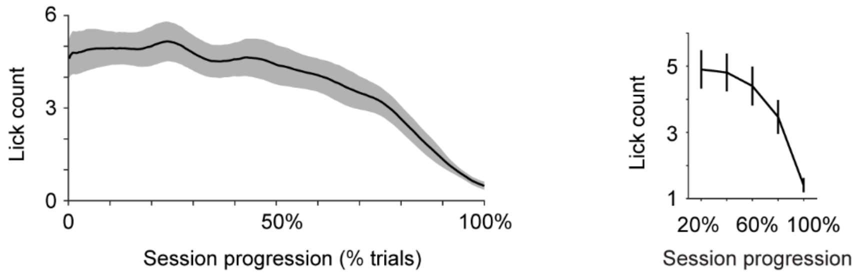


**Figure S2.** Lick count decreased as task progressed within sessions. Left: normalized single-session trajectory of lick count, mean ± s.e.m. Right: averaged every 20% progression. Lick count is defined as the total number of licks emitted in each trial.

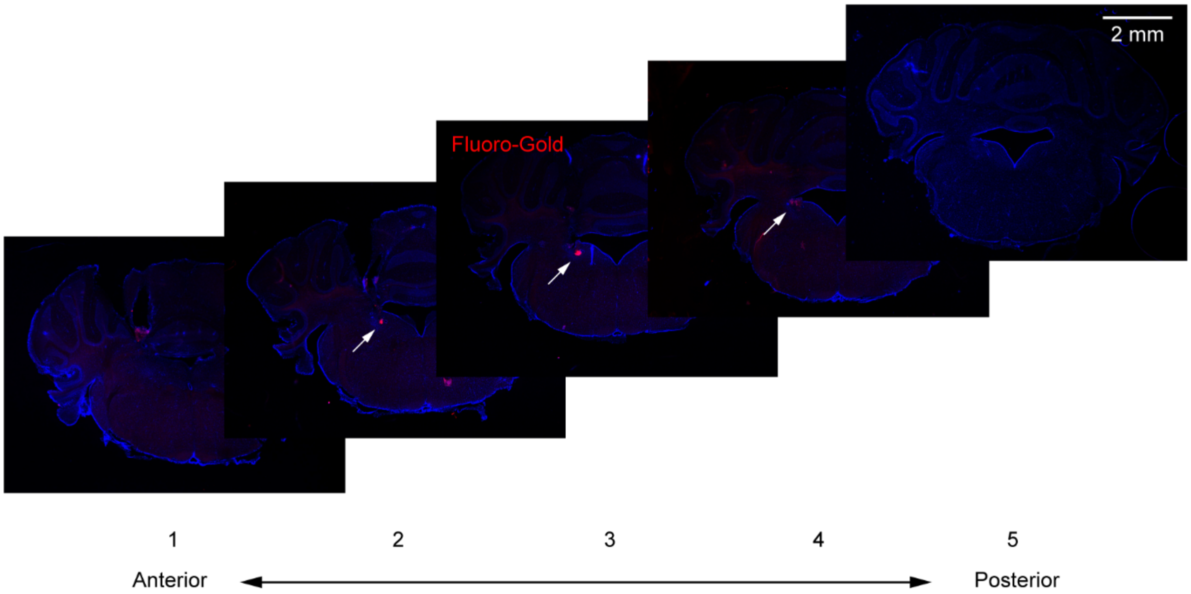


**Figure S3.** 5 consecutive 100 μm coronal sections from a mouse where 300 nL Fluoro-Gold was locally infused. Arrows point to visible Fluoro-Gold (red) in sections 2-4. Fluoro-Gold in the most anterior and posterior sections (1 and 5) is very faint.

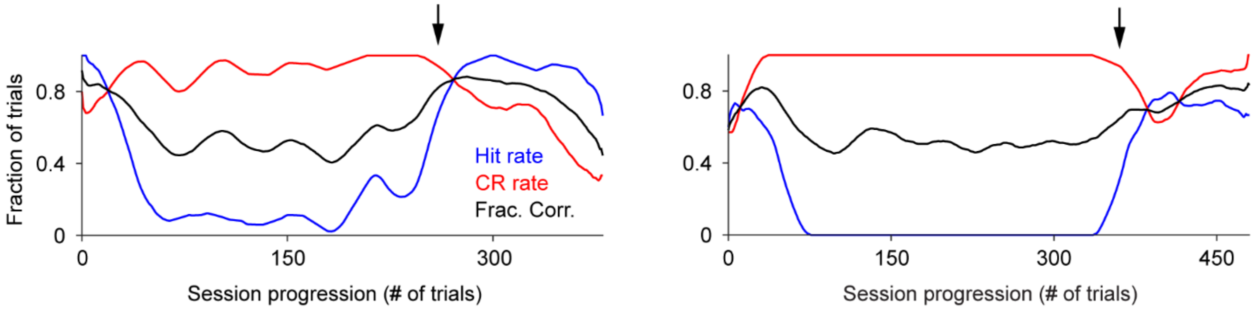


**Figure S4.** Two example behavior sessions of localized clonidine infusion showing that mouse behavior tended to recover later in the session (Hit rate increased and CR rate decreased), indicated by the arrows.

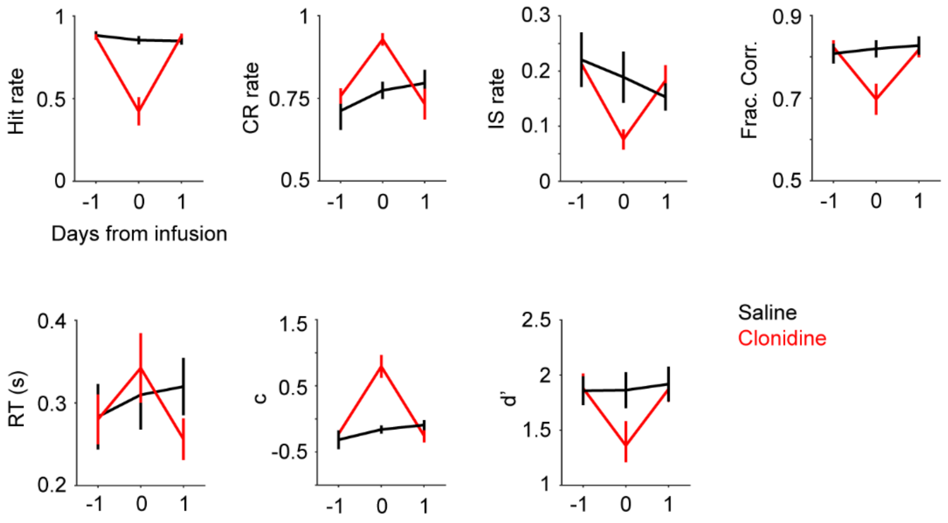


**Figure S5.** Mean Hit rate, CR rate, IS rate, Fraction Correct, RT, decision bias (c) and detection sensitivity (d') for 3 consecutive days (± s.e.m.). Black: saline, n = 6; Red: clonidine, n = 8.

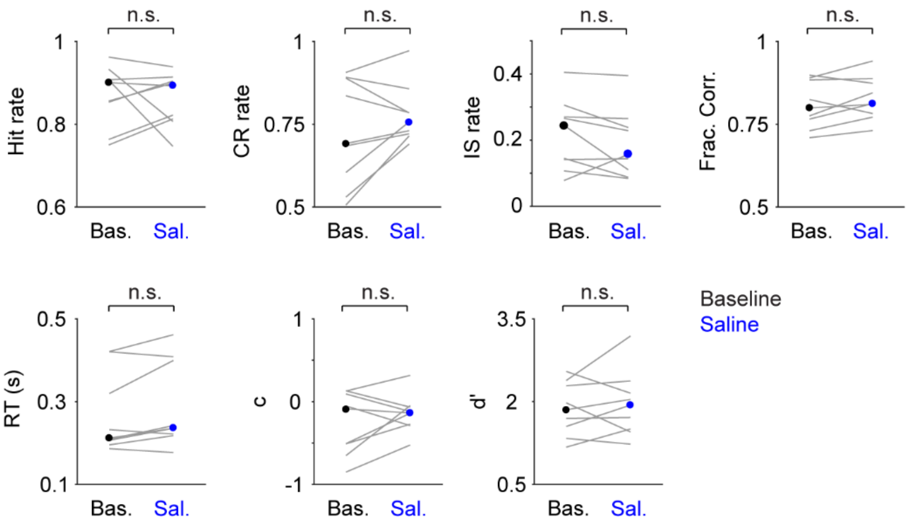


**Figure S6.** Localized saline infusion did not affect behavior. Hit rate, P = 1, Signed rank = 22; CR rate, P = 0.20, Signed rank = 11; IS rate, P = 0.13, Signed rank = 36; Frac. Corr., P = 0.16, Signed rank = 10; RT, P = 0.055, Signed rank = 6; c, P = 0.30, Signed rank = 13; d', P = 0.65, Signed rank = 18. n = 9.

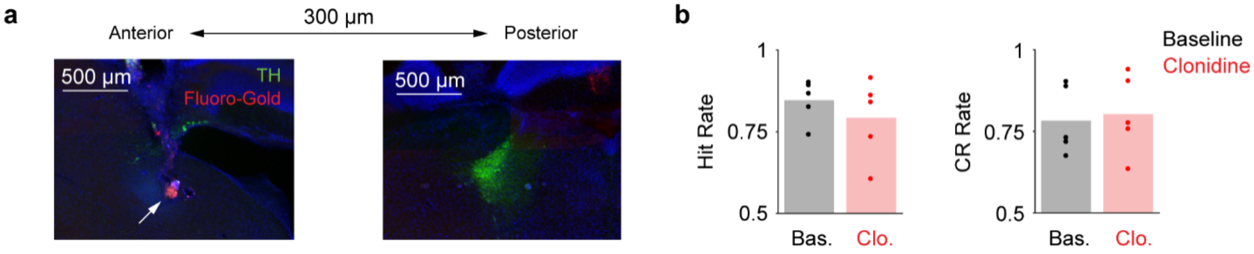


**Figure S7.** Clonidine minimally affected behavior when the infusion was outside of LC. **a.** Example histological sections showing off-target infusion. The location of drug infusion was estimated by Fluoro-Gold (red), which is ~300 μm anterior to the LC (green). **b.** Hit and CR rates during baseline and off-target clonidine sessions (Hit rate, 0.85 ± 0.03 vs. 0.79 ± 0.05; CR rate, 0.78 ± 0.05 vs. 0.80 ± 0.05, mean ± s.e.m., n = 5).

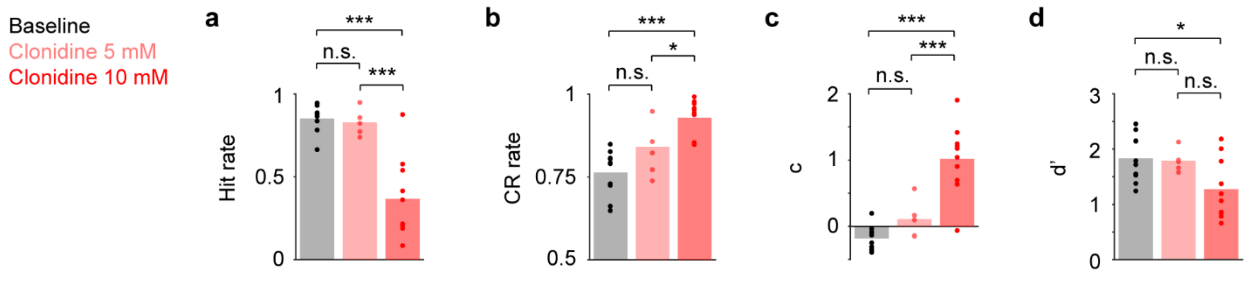


**Figure S8.** Hit rate, CR rate, decision bias (c) and detection sensitivity (d') for baseline (black, n = 10), 5 mM clonidine (light red, n = 5), and 10 mM clonidine (red, n = 10) sessions. **a.** Hit rate: one-way ANOVA, F(2,22) = 23.97, P = 3.0e-6. Baseline vs. 5 mM, P = 0.70; Baseline vs. 10 mM, P = 3.5e-6; 5 mM vs. 10 mM, P = 4.0e-4. **b.** CR rate: one-way ANOVA, F(2,22) = 18.69, P = 1.8e-5. Baseline vs. 5 mM, P = 0.18; Baseline vs. 10 mM, P = 1.2e-5; 5 mM vs. 10 mM, P = 0.013. **c.** Decision bias: one-way ANOVA, F(2,22) = 26.47, P = 1.4e-6. Baseline vs. 5 mM, P = 0.36; Baseline vs. 10 mM, P = 1.2e-6; 5 mM vs. 10 mM, P = 6.6e-4. **d.** Detection sensitivity: one-way ANOVA, F(2,22) = 4.39, P = 0.025. Baseline vs. 5 mM, P = 0.98; Baseline vs. 10 mM, P = 0.029; 5 mM vs. 10 mM, P = 0.12. All post-hoc pairwise tests were Tukey-Kramer.

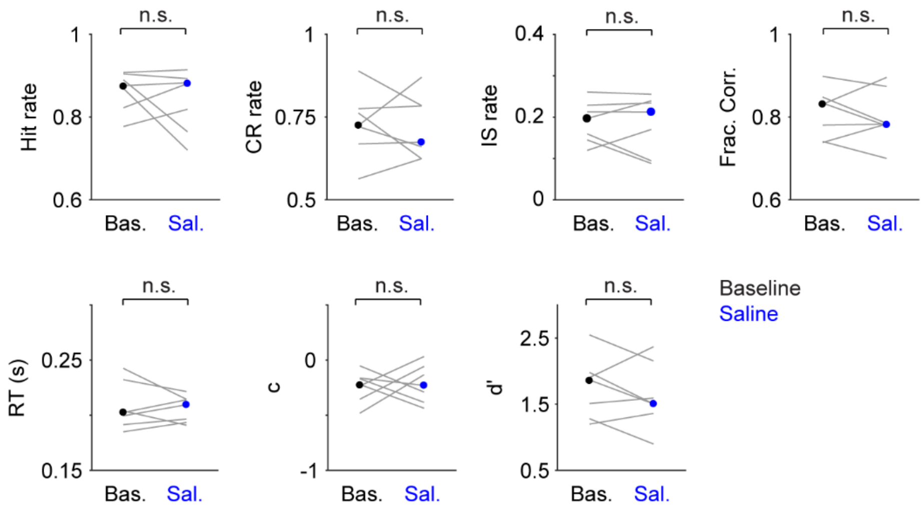


**Figure S9.** Systemic saline administration did not affect behavior. Hit rate, P = 0.81, Signed rank = 16; CR rate, P = 1, Signed rank = 14; IS rate, P = 0.70, Signed rank = 16.5; Frac. Corr., P = 0.81, Signed rank = 16; RT, P = 0.69, Signed rank = 17; c, P = 0.58, Signed rank = 10; d', P = 0.47, Signed rank = 19. n = 7.

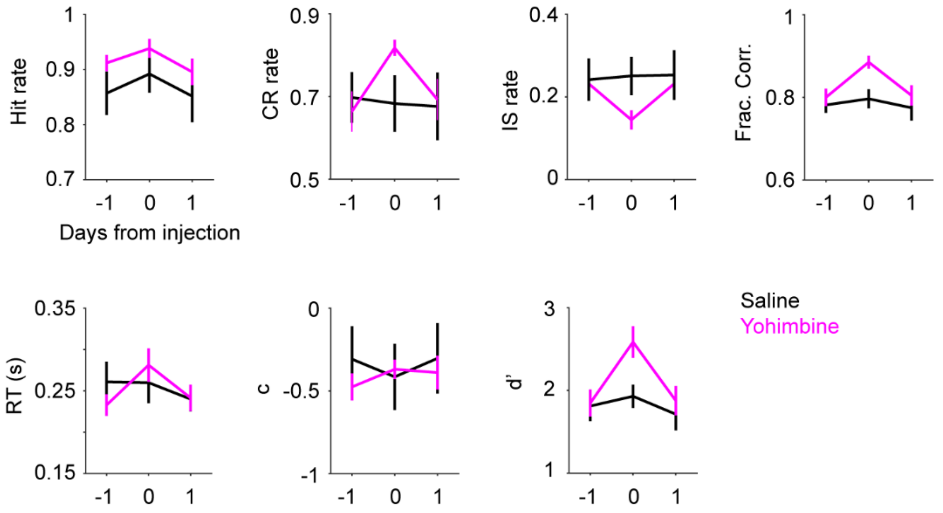


**Figure S10.** Mean Hit rate, CR rate, IS rate, Fraction Correct, RT, decision bias (c) and detection sensitivity (d') for 3 consecutive days (± s.e.m.). Black: saline, n = 7; Magenta: yohimbine, n = 9.

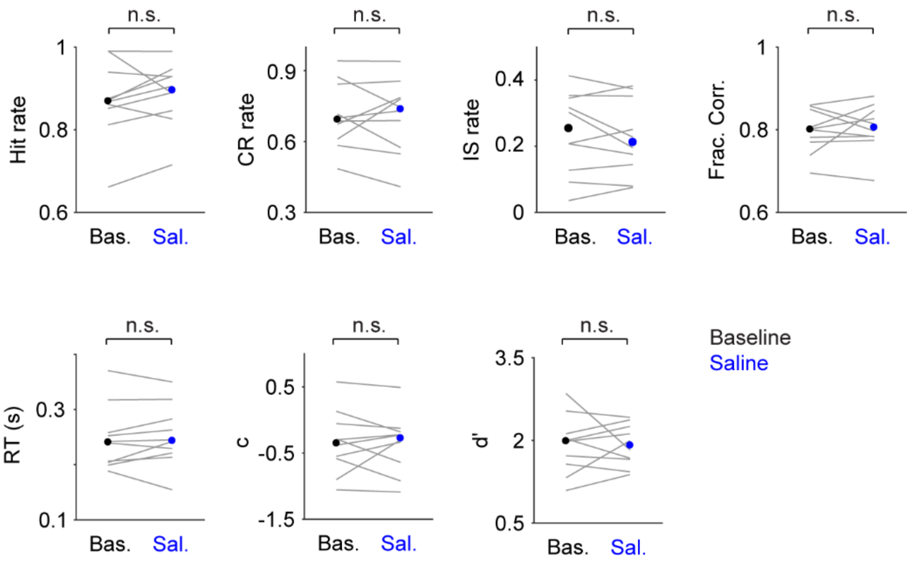


**Figure S11.** Localized saline injection did not affect behavior. Hit rate, P = 0.32, Signed rank = 17; CR rate, P = 0.92, Signed rank = 29; IS rate, P = 0.70, Signed rank = 32; Frac. Corr., P = 0.63, Signed rank = 22; RT, P = 0.43, Signed rank = 19; c, P = 0.77, Signed rank = 31; d', P = 0.86, Signed rank = 25. n = 10.

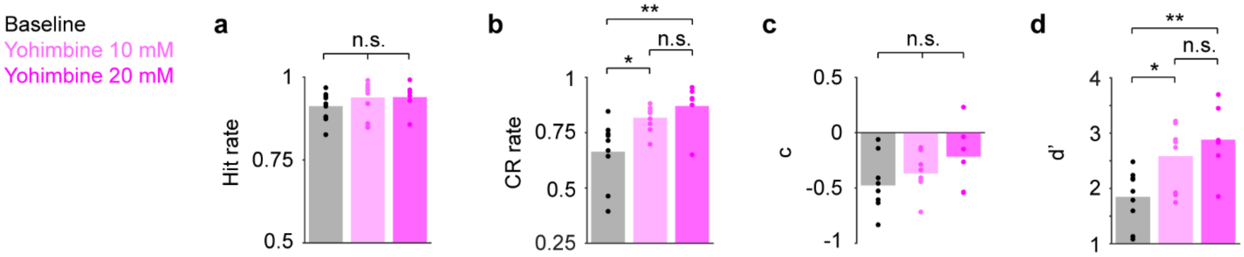


**Figure S12.** Hit rate, CR rate, decision bias (c) and detection sensitivity (d') for baseline (black, n = 9), 10 mM yohimbine (light magenta, n = 9), and 20 mM yohimbine (magenta, n = 6) sessions. **a.** Hit rate: one-way ANOVA, F(2,21) = 0.89, P = 0.42. Baseline vs. 10 mM, P = 0.48; Baseline vs. 20 mM, P = 0.53; 10 mM vs. 20 mM, P = 1.0. **b.** CR rate: one-way ANOVA, F(2,21) = 7.36, P = 0.0038. Baseline vs. 10 mM, P = 0.021; Baseline vs. 20 mM, P = 0.0055; 10 mM vs. 20 mM, P = 0.66. **c.** Decision bias: one-way ANOVA, F(2,21) = 2.17, P = 0.14. Baseline vs. 10 mM, P = 0.61; Baseline vs. 20 mM, P = 0.18; 10 mM vs. 20 mM, P = 0.45. **d.** Detection sensitivity: one-way ANOVA, F(2,21) = 6.9, P = 0.0047. Baseline vs. 10 mM, P = 0.029; Baseline vs. 20 mM, P = 0.0061; 10 mM vs. 20 mM, P = 0.58. All post-hoc pairwise tests were Tukey-Kramer.

| Saline | | | | |
|---|---|---|---|---|
| Mouse number | 1 | | 2 | |
| | Left | Right | Left | Right |
| TH positive cells | 453 | 442 | 586 | 577 |
| TH/c-fos double positive cells | 40 | 41 | 117 | 132 |
| c-fos expression level (%) | 8.8 | 9.3 | 20.0 | 22.9 |
| | P = 0.41 | | P = 0.051 | |

**Table S1.** Quantification of c-fos expression to examine the effect of localized saline infusion on LC activity in 2 awake mice. Saline was infused in the left LC. The right LC serves as a basal level control. Permutation test was performed ($10^5$ iterations) to compare c-fos expression levels between the left and right LC in individual mice.
